# A Plasmid-Derived Machinery Drives Chromosomal Replication Initiation in the Colorectal Cancer-Associated Bacterium *Fusobacterium nucleatum* and Its Close Relatives

**DOI:** 10.64898/2026.10.08.757545

**Authors:** Sophie Pasek, Pierre Brézellec

## Abstract

Chromosome replication initiation in bacterial genomes typically occurs at a single origin, called oriC, through the binding of the DnaA initiator protein to specific chromosomal repeats known as DnaA-boxes. Here, relying on comparative genomics, we show that the family Fusobacteriaceae, which belongs to the order Fusobacteriales alongside the family Leptotrichiaceae, deviates from this paradigm. Our analysis revealed a stark contrast between the two lineages: while *dnaA* flanks oriC in Leptotrichiaceae, it is notably missing from Fusobacteriaceae genomes. Adjacent to oriC in Fusobacteriaceae, we identified a gene encoding replication initiator protein A (RPA). We characterized RPA as a Rep-type initiator involved in plasmid replication that likely originated from an iteron-containing plasmid, iterons being the plasmid analogues of DnaA-boxes. Phylogenetic reconstruction indicates that the gene encoding RPA was acquired together with its associated iterons and subsequently domesticated by the Fusobacteriaceae ancestor, successfully displacing the ancestral chromosomal *dnaA* gene at the origin of replication. This shift to a plasmid-derived replication system may represent an adaptive trait of the Fusobacteriaceae, potentially offering new insights into the biology of *Fusobacterium nucleatum*, a member of this family known to be a major driver of colorectal cancer progression within the tumor microenvironment.

## Introduction

The replication of bacterial chromosomes is highly regulated, beginning at a specific region called the origin of replication (oriC). In this context, the DnaA protein acts as the master regulator, recognizing and binding to specific DNA sequences known as DnaA-boxes to form a nucleoprotein filament. This assembly generates the mechanical tension required to trigger the opening of the DNA unwinding element (DUE), an AT-rich region adjacent to the binding sites. This initial melting of the double helix is a prerequisite for the loading of replicative helicases, which then drive DNA synthesis in opposite directions (Leonard & Grimwade, 2015). Exceptions to this paradigm have been documented for some endosymbionts with reduced genomes, such as *Blochmannia floridanus*, whose genome lacks the *dnaA* gene (Gil *et al*., 2003). However, the most numerous documented violations of the DnaA-oriC-dependent replication initiation mechanism are found in the cyanobacteria. For example, the free-living cyanobacteria *Cyanobacterium aponinum* PCC 10605 and *Geminocystis s*p. NIES-3708 do not encode *dnaA* and replicate their chromosomes from multiple origins (Ohbayashi *et al*., 2020). *Synechocystis sp*. PCC 6803 and *Nostoc sp*. PCC 7120 represent another case. Their chromosomes are replicated asynchronously from multiple origins, and while they host a *dnaA* gene, deletion of this gene affects neither the growth nor the chromosomal replication activity of these species, suggesting that they may replicate their chromosomes through an unknown DnaA-oriC-independent mechanism (Ohbayashi *et al*., 2016).

The involvement of *Fusobacterium nucleatum* in colorectal cancer (Kostic *et al*., 2012) has fueled significant scientific interest in this bacterium (Udayasuryan *et al*., 2024). In a recent study (Molteni *et al*., 2024), the genetic diversity and phylogenetic relationships of species within the genus *Fusobacterium* were assessed. To conduct their analysis, the authors identified 120 core genes present in more than 300 *Fusobacterium* genomes. Among these, they retained only genes longer than 1000 bp, resulting in a set of 45 core genes. Upon examining this dataset, we observed the absence of the *dnaA* gene, which is typically ∼1500 bp long, suggesting that it is not conserved within this genus. Interestingly, consistent with this result, the authors who provided the first genome sequence of *Fusobacterium nucleatum* subsp. *nucleatum* were unable to find *dnaA* (Kapatral *et al*., 2002). They tentatively identified an ORF of 637 amino acids, located near the *recF* gene, as encoding the DnaA protein. However, the length of this ORF is unusual for a DnaA protein; for instance, *Escherichia coli* DnaA is only 467 amino acids long. In addition, its Pfam domain architecture (Blum *et* al., 2025) deviates from that of a typical DnaA protein: it is composed of a single PF10134 domain (Replication initiator protein A / RPA), whereas a classic DnaA protein comprises three domains, namely PF11638 (DnaA N-terminal domain), PF00308 (Bacterial DnaA ATPase domain), and PF08299 (Bacterial DnaA protein helix-turn-helix). These findings led us to re-evaluate the DnaA-oriC-dependent replication initiation mechanism in the order Fusobacteriales, to which the genus *Fusobacterium* belongs.

Through a comparative genomics analysis, we demonstrate herein that the ancestor of the Fusobacteriaceae family domesticated a plasmid-derived gene encoding a Rep-type initiator, which replaced the ancestral chromosomal *dnaA* gene at its original locus. This represents the first documented shift from a chromosomal to a plasmid-derived initiator across Bacteria. We discuss the implications of this functional replacement and its impact on the biology of this family, which includes *Fusobacterium nucleatum*, a key driver of colorectal cancer progression.

## Materials and methods

### Databases

We relied on DoriC 12.0 (Dong, Luo, and Gao, 2023) to locate both the oriC and *dnaA* genes. This database identifies replication origins by combining indicator genes (e.g., *dnaA, dnaN*) with Z-curves. These curves capture the compositional asymmetries between the leading and lagging strands. Such asymmetry stems from the distinct synthesis mechanisms of each strand during DNA replication, which exposes them to different mutational pressures, leading to an enrichment of guanine on one strand and cytosine on the other (Lobry and Sueoka, 2002).

To perform the visual inspection of these profiles, we used SkewDB (Hubert, 2022), which provides GC- and TA-skews for over 53,000 chromosomes and plasmids via a user-friendly interface.

### Dataset

To study the distribution of the chromosomal replication initiator gene *dnaA* and the oriC region within the order Fusobacteriales, we built a dataset by querying DoriC 12.0, specifying ‘Fusobacteriales’ in the ‘Lineage’ field. The search returned 160 records, many of which corresponded to multiple sequencing entries for the same species or strains. From these, we discarded one record for which DoriC results were inaccessible (NCBI RefSeq assembly GCF_002573475.1), along with 12 records corresponding to genome assemblies suppressed by NCBI for various reasons (e.g., “unverified source organism” or “contamination”). This curation step resulted in a final dataset comprising 37 species, 3 subspecies, and 52 strains (see Supplementary Material Table S1).

Although DoriC 12.0 indexes a vast number of bacterial genomes, it occasionally misses biologically relevant species. To expand our coverage of the Fusobacteriales, we compiled an additional dataset comprising species classified as “Reference Proteomes” by UniProt (UniProt Consortium, 2025). These proteomes notably include well-studied model organisms and other species of significant interest for biomedical research. To identify these Fusobacteriales species, we queried UniProt Proteomes for entries matching (taxonomy_id:203491) AND (proteome_type:1), where 203491 is the TaxID for Fusobacteriales and proteome_type:1 refers to “Reference Proteomes” (UniProt Proteomes release 2025). Twenty-two Fusobacteriales species indexed in UniProt Proteomes matched these criteria (see Supplementary Material Table S2), five of which were not recorded in DoriC 12.0. Given that our subsequent analysis requires locating oriC, we narrowed the selection to species available in SkewDB. This selection process left us with the following three species: *Fusobacterium nucleatum subsp. nucleatum* (strain ATCC 25586), *Ilyobacter polytropus* (strain ATCC 51220), and *Fusobacterium hominis*. These three species belong to the Fusobacteriaceae family.

To sum up, our dataset comprises 40 species (37 retrieved from DoriC and 3 from UniProt) and 3 subspecies (collected from DoriC). However, DoriC contains genomes at varying stages of assembly (from contigs and scaffolds to complete genomes). For this reason, we focused part of our analysis exclusively on the subset of 23 species whose genomes are complete (20 species collected from DoriC and 3 from UniProt), see Supplementary Material Table S1 (the symbol “v” denotes records with plotted GC-skew).

### Softwares

We used the HHpred tool via its web server (https://toolkit.tuebingen.mpg.de/tools/hhpred). This tool enables sensitive protein domain and structure prediction by utilizing pairwise alignment of Hidden Markov Models (HMMs) to detect remote sequence homologies (Gabler *et al*., 2020).

We used MEME (https://meme-suite.org/meme/tools/meme) to search for iterons (Bailey & Elkan, 1994). Iterons are DNA sequences within replication origins of plasmids where initiator proteins bind to trigger replication initiation. MEME identifies novel, ungapped motifs (recurring, fixed-length patterns) in a set of sequences, which in our case were the intergenic regions containing oriC.

Custom Python and R scripts were written to compute and plot GC skews for Figure 1 (see file CODES.tgz in the Zenodo repository https://zenodo.org/records/23887865).

**Figure 1.**
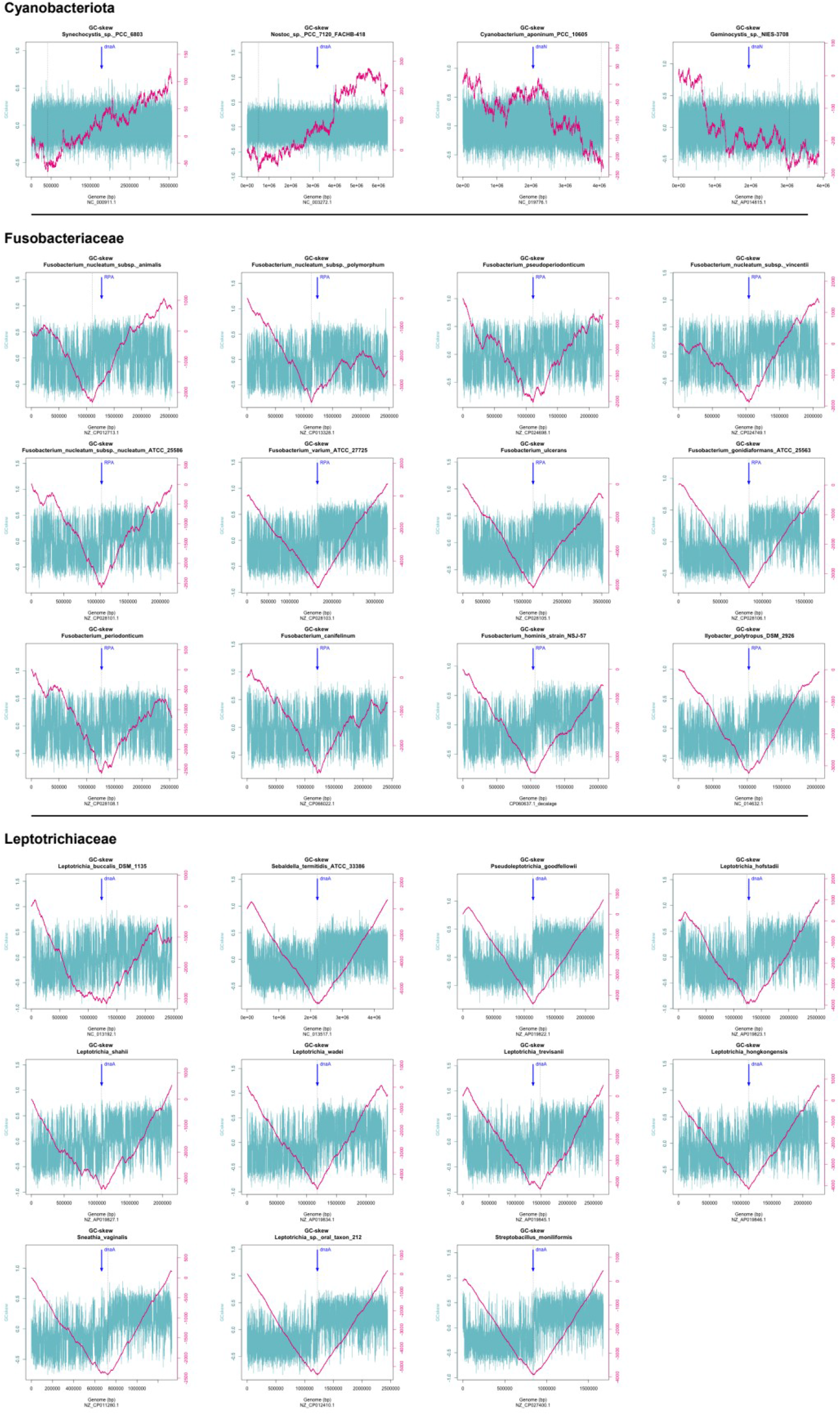
GC-Skew analysis supports single-oriC replication in Fusobacteriales. The bacterial replication origin can be localized *in silico* through a GC skew curve, which characteristically has a V-shaped profile. The inflection point (the bottom of the ‘V’) shows the transition between the leading and lagging strands. Some cyanobacteria do not follow the canonical, *dnaA*-dependent, single-oriC replication initiation mechanism, meaning their curves lack a clear inflection point (small top section of the image). In contrast, members of Fusobacteriaceae (middle section) and Leptotrichiaceae (bottom sectoin) exhibit V-curves, strongly suggesting they initiate replication from a unique oriC. The blue arrow indicates the location of *dnaN* in cyanobacteria (where *dnaA* is not always conserved), *dnaA* in Leptotrichiaceae, and the gene encoding replication protein A (RPA) in Fusobacteriaceae. Interestingly, in both Fusobacteriaceae and Leptotrichiaceae, the oriC locus closely matches that of the RPA-encoding gene and *dnaA*, respectively.

## Results

### GC-skew profiles of Fusobacteriales species support a single oriC-dependent replication initiation model

The cumulative GC-skew of a bacterial genome typically forms a distinct V-shaped pattern. The inflection point at the bottom of this ‘V’ marks the transition between the leading and lagging strands and is commonly used to predict the chromosomal replication origin (Lobry and Sueoka, 2002).

We investigated the GC-skew patterns of the order Fusobacteriales, which comprises two main families: Leptotrichiaceae and Fusobacteriaceae. We visually inspected their profiles using SkewDB, focusing on the 23 completely sequenced genomes within our dataset (see Dataset section and Supplementary Material Table S1). This revealed that all of them exhibit V-shaped profiles (see Figure 1, where we also included four cyanobacterial species known to violate the single-oriC replication initiation mechanism to clearly highlight the contrast between V-shaped and non-V-shaped profiles).

Our results thus strongly suggest that replication initiation in Fusobacteriales species follows a single-oriC-dependent replication initiation model.

### Species within the Leptotrichiaceae family harbor the chromosomal replication initiator gene *dnaA* located next to oriC, whereas Fusobacteriaceae species lack the *dnaA* gene

As mentioned in the introduction, evidence suggests that the genus *Fusobacterium*, which belongs to the Fusobacteriaceae family, lacks the *dnaA* gene. We thus decided to investigate the distribution of *dnaA* within the order Fusobacteriales. To achieve this, we examined the DoriC 12.0 annotations of the genomes in our dataset, as this database records, among other information, the location of *dnaA* whenever it is present (see Supplementary Material Table S1).

We observed that all Leptotrichiaceae species harbor a *dnaA* gene located next to oriC, whereas nearly all Fusobacteriaceae lack *dnaA*. The only exceptions to this rule were three Fusobacteriaceae species where a gene annotated as *dnaA* was found adjacent to oriC. Specifically, this was the case for two different releases of *Fusobacterium perfoetens* (*F. perfoetens* and *F. perfoetens* ATCC 29250, encoding proteins WP_035945371 and WP_176828875, respectively), one release of *Fusobacterium sp*. (*F. sp*. FSA-380-WT-3A, encoding protein WP_370456854), and the single release of *Hypnocyclicus thermotrophus* (encoding protein WP_134113243). These exceptions prompted us to perform a more detailed analysis of the four proteins encoded by these *dnaA*-annotated genes.

Our analysis revealed that these four protein-coding genes were approximately 600 aa long. Given that bona fide DnaA proteins are typically around 500 aa long, this 100-aa discrepancy led us to re-annotate them using HHpred (Gabler *et al*., 2020). Searches were performed against the Pfam-A v37.4 database (Blum *et al*., 2025) using these proteins as queries. HHpred identified the N-terminal region in all four cases as a Replication initiator protein A domain (Pfam PF10134/RPA) with 99.9% confidence (see file HHPRED.zip in the Zenodo repository https://zenodo.org/records/23887865). Since the typical Pfam domain architecture of a canonical DnaA protein consists of three specific domains, namely PF11638 (DnaA_N), PF00308 (DnaA_ATPase), and PF08299 (DnaA_C), we concluded that these four genes had been misannotated and actually encode Replication initiator protein A.

We then focused on the three Fusobacteriaceae species corresponding to UniProt reference proteomes that were not included in DoriC 12.0 (see the Dataset section). Since no instance of a *dnaA* gene was found in the Fusobacteriaceae species examined so far, we investigated whether these three species possess the *dnaA* gene by searching their proteomes using BLASTP for homologs of the DnaA protein from *Sebaldella termitidis* (a member of the Leptotrichiaceae family). This search yielded no significant hits (E-value < 10^-6^), suggesting the absence of *dnaA* in these genomes (see file BLAST.txt in the Zenodo repository https://zenodo.org/records/23887865).

Overall, our analysis strongly suggests that while Leptotrichiaceae species possess the chromosomal replication initiator gene *dnaA* located next to oriC, this gene is absent in members of the Fusobacteriaceae family. Note that in some species, such as *Leptotrichia wadei*, DoriC predicts two oriC loci flanking *dnaA* (*i*.*e*., an oriC on each side of the gene) within the same genome release. To the best of our knowledge, DoriC does not predict bipartite oriC loci (see Leonard & Grimwade, 2015, for examples of a bipartite oriC configuration); these are, therefore, two different predicted locations for a single oriC. However, such ambiguities do not undermine our main conclusion.

### A gene encoding a replication initiator protein A (RPA) is adjacent to *oriC* in Fusobacteriaceae

As the replication initiation gene *dnaA* often resides near oriC, we hypothesized that a new initiation factor might be located nearby in species of the Fusobacteriaceae family. Therefore, we investigated the genomic context of this region, defined here as the three loci flanking each side of oriC, across the Fusobacteriaceae of our dataset.

We first examined the oriC regions of the species annotated by DORIC 12.0. We found that all these Fusobacteriaceae species carry a gene encoding a ‘Replication initiator protein A’ (∼610 aa) located next to oriC (see Supplementary Material Table S1).

We then focused on the three species not stored in DORIC 12.0 that are associated with a UniProt reference proteome and recorded in SkewDB (see section Dataset), namely: *Fusobacterium nucleatum subsp. nucleatum* (strain ATCC 25586), *Ilyobacter polytropus* (strain ATCC 51220), and *Fusobacterium hominis*. We used SkewDB to predict the oriC locations for these three species.

For *Fusobacterium hominis* (accession number CP060637; genome length: 2,072,767 bp), we located the oriC region at approximately position 1,600,000 bp. Since this genome is indexed in KEGG (Kanehisa & Goto, 2000), we used this platform to examine this genomic region. At positions 1,604,033–1,605,823, we identified a gene (locus tag H9Q81_07835) encoding a 596-amino-acid protein. This protein is annotated as a ‘Replication initiator protein A’ and carries the PF10134/RPA Pfam domain (*i*.*e*., replication initiator protein A).

For *Ilyobacter polytropus* (accession number NC_014632.1; genome length: 2,046,464 bp), we located the oriC region near position 0. Since this assembly is not available in KEGG, we visualized the region using the “Graphics” utility of the NCBI Nucleotide database. The gene located at positions 24–1,862 (locus tag ILYOP_RS00010) encodes a 612-aa protein annotated as a ‘Replication initiator protein A’.

Finally, we examined *Fusobacterium nucleatum* subsp. *nucleatum* (accession number NZ_CP028101.1; genome length: 2,180,101 bp), for which we located the oriC region around position 425,000 bp. This region was also visualized using the “Graphics” utility of the NCBI Nucleotide database. A gene (locus tag C7Y58_RS02225) located at positions 424,800–426,713 was found to encode a 612-aa protein, which is likewise annotated as a ‘Replication initiator protein A’.

Overall, our analysis demonstrates that a gene encoding a Replication initiator protein A is adjacent to *oriC* in the Fusobacteriaceae.

### Replication initiator protein A is involved in the replication initiation of plasmids and likely originated from the replication initiator gene of an iteron-containing plasmid

While establishing that Leptotrichiaceae species harbor the chromosomal replication initiator gene *dnaA* whereas Fusobacteriaceae lack it, we analyzed misannotated DnaA proteins from Fusobacteriaceae species using Hhpred, which enables sensitive protein domain prediction. This analysis identified the Pfam domain PF10134/RPA (Replication initiator protein A domain) in their N-terminal region with 99.9% confidence. Interestingly, these N-terminal regions also exhibited significant homology to six other Pfam domains, all identified with >98% confidence (see file HHPRED.zip in the Zenodo repository https://zenodo.org/records/23887865). A brief summary of the InterPro annotations for these domains is provided in the table below:

**Table 1.** Each row represents a Pfam domain matching with high confidence the N-terminal region of Fusobacteriaceae proteins misannotated as DnaA. The ‘Pfam domain’ column indicates the name of the Pfam domain, and ‘InterPro family’ indicates the InterPro entry into which the Pfam domain has been integrated. The ‘Description’ column compiles information extracted from InterPro domain annotations.

| Pfam domain | InterPro family | Description (derived from InterPro database) |
|---|---|---|
| PF07112 - DUF1368 | IPR010778 | Includes proteins seemingly specific to red algae plasmids; function remains unknown |
| PF07042 - TrfA | IPR010751 | The *trfA* operon of broad-host-range IncP plasmids is essential for activating the vegetative origin of replication across diverse species |
| PF11826 - RctB_central | IPR021781 | Harbored by the replication initiator protein RctB, encoded by a gene located on *Vibrio cholerae* chromosome 2 (likely of plasmid origin (Jha *et al*., 2017)), as well as in related sequences, predominantly within Vibrionales |
| PF06504 - RepC | IPR010522 | Covers various bacterial replication protein C (RepC) sequences |
| PF04796 - RepA_C | IPR006881 | Comprises proteins involved in plasmid replication. |
| PF01051 - Rep3_N | IPR000525 | Represents the winged helix 1 (WH1) domain found in plasmid replication initiator proteins (e.g., *E. coli* RepA, which binds DNA repeats flanking repA). |

In summary, most of these domains are associated with plasmid proteins involved in replication initiation, including PF07042/TrfA, PF11826/RctB_central, and PF01051/Rep3_N. Interestingly, PF07042/TrfA and PF01051/Rep3_N belong to initiator proteins that trigger replication by binding to iterons (short, repeated DNA sequences) (Konieczny & Helinski, 1997; Giraldo & Fernández-Tresguerres, 2004). The PF11826/RctB_central domain is found in the *Vibrio cholerae* RctB protein; the binding of RctB to an array of six 12-mer sites (iterons) within the chromosome 2 (Chr2) origin of *Vibrio cholerae* is essential for the replication initiation of this replicon (Chatterjee *et al*., 2020). It should be noted that Chr2 of *V. cholerae* is hypothesized to have evolved from an iteron-family plasmid (Chatterjee *et al*., 2020; see also Adam *et al*., 2022).

Overall, this strongly suggests that the Replication initiator protein A is involved in the replication initiation of plasmids and likely originated from the replication initiator gene of an iteron-containing plasmid.

### In the Fusobacteriaceae family, the gene encoding Replication initiator protein A (RPA) replaced the *dnaA* gene at its ancestral locus

A comparison of the oriC genomic context between *Sebaldella termitidis* (Leptotrichiaceae) and *Fusobacterium nucleatum subsp. nucleatum* (Fusobacteriaceae) revealed a highly conserved gene order: *recF/yaaA/dnaA*–oriC–*rpmH/rnpA/yidD* versus r*ecF/yaaA/*RPA–oriC– *rpmH/rnpA/yidD*, respectively (see Supplementary material Table S1). They differ exclusively in the gene upstream of oriC, where a gene encoding Replication initiator protein A (RPA) occupies the position of *dnaA* in *Fusobacterium nucleatum subsp. Nucleatum*. This raises the question of whether a replacement of *dnaA* by a gene encoding RPA occurred at this specific locus in Fusobacteriaceae, whereas Leptotrichiaceae retained the ancestral arrangement. To trace the evolutionary history of this locus (see below), we used parsimony to reconstruct the ancestral oriC context of Fusobacteriales, based on the 15 Fusobacteria species from our dataset that have complete genomes and are represented in the most comprehensive Fusobacteria phylogeny currently available (Gupta & Sethi, 2014; see Figure 2).

**Figure 2.**
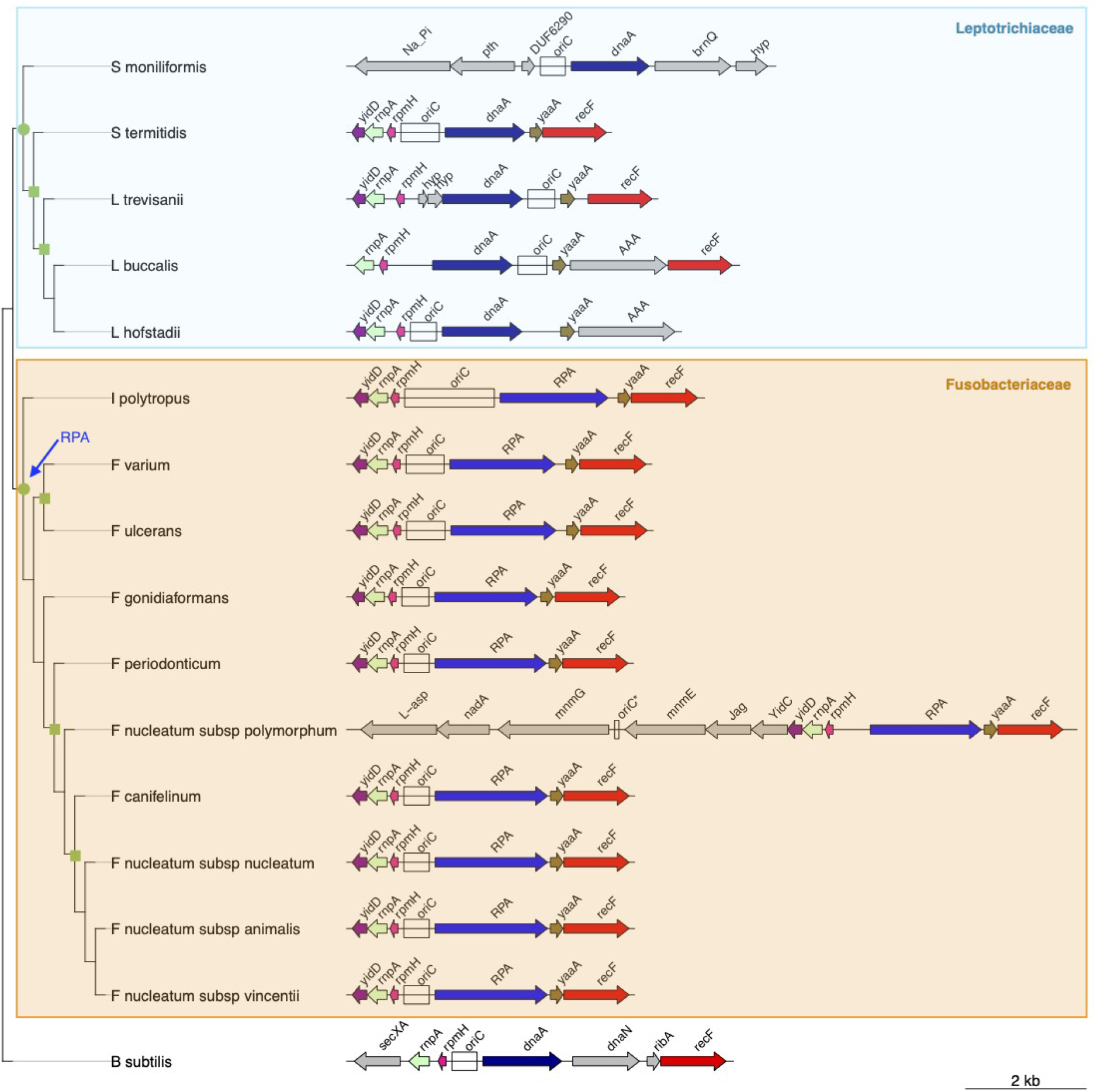
The tree shown in the figure maps a subset of our species dataset (restricted to fully sequenced species) onto a schematic phylogenetic tree of the order Fusobacteriales, adapted from Gupta & Sethi (2014). *Bacillus subtilis* is used as a deeply branching outgroup species. On nodes, green circles and squares indicate bootstrap values of 100% and at least 70%, respectively. Each species is associated with the genomic context of its oriC, defined here as the three flanking genes on each side. Arrows represent genes, the abbreviation “hyp” indicates a hypothetical protein, and oriC region is represented by an empty rectangle. Parsimony was used to reconstruct the history of the oriC genomic context. We inferred that *recF*/*yaaA*/RPA – oriC – *rpmH*/*rnpA*/*yidD* was the ancestral state for Fusobacteriaceae, and *recF*/*yaaA*/*dnaA* – oriC – *rpmH*/*rnpA*/*yidD* for Leptotrichiaceae. This strongly suggests that the gene encoding RPA replaced *dnaA* at its ancestral site at the root of the Fusobacteriaceae family; this event is highlighted by a blue arrow pointing to the node representing the root of this clade. Note that the atypical oriC location in *F. nucleatum subsp. polymorphum* (marked by an asterisk) likely stems from an annotation error.”

The *recF*/*yaaA*/RPA–oriC–*rpmH*/*rnpA*/*yidD* cluster is clearly the ancestral oriC genomic context of the Fusobacteriaceae family. The atypical oriC location in *Fusobacterium nucleatum subsp. polymorphum* (marked by an asterisk in Figure 2) likely stems from an annotation error, as this complete ancestral cluster is found just three genes downstream of the predicted oriC.

Within the Leptotrichiaceae family, the oriC genomic context of *Sebaldella moniliformis* is highly divergent, sharing no common features with other species in this group except for the presence of *dnaA* next to oriC. Interestingly, the oriC genomic context of *Sebaldella termitidis* mirrors the ancestral state of the Fusobacteriaceae family, with the exception of *dnaA*, which occupies the position of the gene encoding RPA. The remaining species (*Leptotrichia trevisanii*,

*L. buccalis*, and *L. hofstadii*) share a core set of genes: *dnaA, yaaA, rpmH*, and *rnpA*. Beyond this common core, *L. trevisanii* shares *recF* with *L. buccalis*, and *yidD* with *L. hofstadii*. All these genes, except *dnaA*, are embedded in the oriC genomic context of the ancestor of the Fusobacteriaceae family. Taken together, this strongly suggests that the ancestral state for Leptotrichiaceae is *recF*/*yaaA*/*dnaA*–oriC–*rpmH*/*rnpA*/*yidD*. We thus conclude that the gene encoding RPA replaced *dnaA* at its ancestral site at the root of the Fusobacteriaceae family.

This result is consistent with other evidence. Recently, a new species was identified as a member of the order Fusobacteriales (Miyazaki *et al*., 2023). Based on its distinctive characteristics, the authors proposed a novel family, genus, and species within this order, named Haliovirgaceae fam. nov., *Haliorigas* gen. nov., and *Haliovirga abyssi* sp. nov. They showed that Fusobacteriaceae and Leptotrichiaceae are more closely related to each other than to

Haliovirgaceae, meaning that Fusobacteriaceae and Leptotrichiaceae are sister families, with Haliovirgaceae branching as an outgroup. Using SkewDB, we identified the oriC region of this species and established its genomic context (see file Haliovirga.txt in the Zenodo repository https://zenodo.org/records/23887865) as *yidD*–*rnpA*–*rpmH*–oriC–*dnaA*–*yaaA*–*recF*. This suggests that the ancestor of the Fusobacteriales harbored a *dnaA* gene located next to oriC. While these traits were inherited by the common ancestor of all families, *dnaA* was subsequently replaced by a gene encoding RPA within the Fusobacteriaceae lineage.

Overall, our results strongly suggest that, within the Fusobacteriaceae, the gene encoding replication protein A (RPA) replaced the *dnaA* gene at its ancestral locus.

### The gene encoding Replication initiator protein A (RPA) was likely accompanied by its iterons when it replaced *dnaA* at its ancestral locus

We previously showed that the gene encoding Replication initiator protein A (RPA) is involved in plasmid replication initiation and likely originated from an iteron-containing plasmid (see section “Replication initiator protein A is involved in the replication initiation of plasmids and likely originated from the replication initiator gene of an iteron-containing plasmid”). In such mobile elements, the plasmid-encoded initiator protein binds to iterons (*i*.*e*., repeated DNA sequences) to promote replication initiation (Konieczny *et al*., 2014). Iterons are analogous to DnaA-boxes; the interaction of the initiator with the iterons melts the DNA unwinding element (DUE), leading to DNA strand separation at the origin of replication.

Using MEME (Bailey & Elkan, 1994), we searched for such sites within the oriC regions of the nine Fusobacteriaceae species used in the previous section (*i*.*e*., those with a complete genome represented in the Fusobacteria phylogeny from Gupta & Sethi, 2014). Given that iterons are short sequences (17–22 bp) characterized by variable numbers (typically 3 to 7) and spacing among plasmids (Konieczny *et al*., 2014), we ran MEME using parameters that accommodate these constraints. The “Select the number of motifs” parameter was set to 3, to search for three types of motifs/iterons. The “Select the site distribution” parameter was set to ‘Any number of repetitions’, allowing MEME to assume that each sequence may contain any number of non-overlapping occurrences of each motif. Finally, the motif width was constrained to a range of 15 to 25 nucleotides, while all other settings were left at their default values.

The three motifs discovered by MEME were statistically significant (E-value < 0.05) (see Figure 3 and file MEME.zip in the Zenodo repository https://zenodo.org/records/23887865). The most significant motif (E-value = 10^-104^; width: 21 nucleotides) occurred at least three times within each of the nine selected oriC regions. Conversely, the other two motifs were present only once in most of these regions. Given that a single iteron occurrence is presumably insufficient to trigger open complex formation, and considering that the number of repeats typically ranges from three to seven in documented plasmids, we concluded that the most significant motif likely corresponds to the functional iterons. Although only experimental assays can validate the relevance of this *in silico* study, our result is consistent with the hypothesis that the gene encoding RPA was accompanied by its iterons when it replaced *dnaA* at its ancestral position.

**Figure 3.**
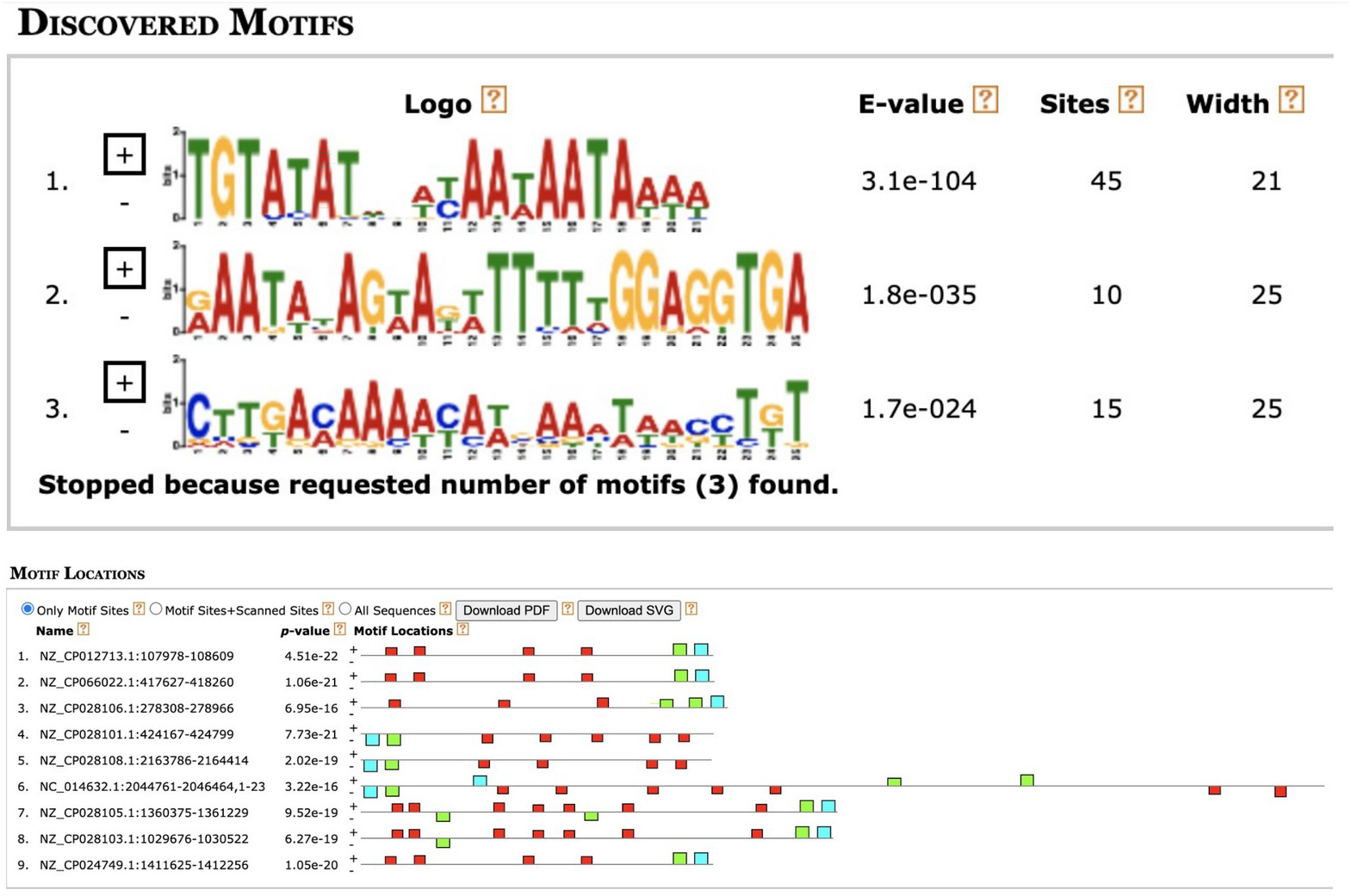
The replacement of *dnaA* at its ancestral locus by the gene encoding replication initiator protein A was likely accompanied by its cognate iterons. Three conserved motifs were identified across nine Fusobacteriaceae oriC regions using MEME (Bailey & Elkan, 1994). The “Discovered Motifs” panel displays the three motifs found by MEME (named 1, 2, and 3 in decreasing order of significance), alongside their E-values, total occurrences, and widths. All three motifs are statistically significant (E-value < 0.05). The “Motif Locations” panel displays the location and number of occurrences of these motifs within each oriC region, with red, blue, and green boxes representing motifs 1, 2, and 3, respectively. Given that a single iteron occurrence is presumably insufficient to trigger open complex formation, and considering that the number of repeats typically ranges from three to seven in known plasmids, the most significant motif (motif 1, red box) is inferred to correspond to the functional iterons.

## Discussion

In most bacteria, chromosome replication is initiated at a single site, oriC, through the binding of the bacterial DnaA initiator protein to “DnaA boxes” (*i*.*e*., short repeated DNA sequences); this binding leads to the melting of the DNA unwinding element (DUE), which promotes open complex formation at the origin (Leonard & Grimwade, 2015). While exceptions to this rule are rare, upon analyzing a dataset of conserved core genes within the genus *Fusobacterium* (Molteni *et al*., 2024), we realized that the *dnaA* gene was missing. This finding prompted us to investigate the validity of the canonical DnaA-single oriC model across the order Fusobacteriales, spanning both the Leptotrichiaceae and Fusobacteriaceae families, the latter including the genus Fusobacterium.

We first demonstrated that Fusobacteriales genomes display V-shaped GC-skews, strongly suggesting a single oriC-dependent replication initiation mechanism (see Figure 1). We then showed that while *dnaA* flanks oriC in Leptotrichiaceae, it is notably missing from the genomes of species belonging to the Fusobacteriaceae family. In species of this family, adjacent to oriC, we identified a gene encoding Replication initiator protein A (RPA), which we characterized as a Rep-type initiator of plasmid origin (see Table). Phylogenetic reconstruction (see Figure 2) indicates that RPA was likely acquired alongside its iterons (see Figure 3) and subsequently domesticated by the Fusobacteriaceae ancestor, successfully displacing the ancestral chromosomal *dnaA* gene at the origin of replication.

The immediate benefit to the cell of replacing the chromosomal replication initiator gene *dnaA* with a plasmid-derived alternative is difficult to perceive. Nonetheless, it can be assumed that this substitution did not perturb the host cell cycle, or did so only minimally. This hypothesis is supported by the fact that *dciA*, the ancestral replicative helicase loader essential for bacterial replication initiation, remains present in the *Fusobacteriaceae* family (Brézellec *et al*., 2016), despite the replacement of *dnaA* by the plasmid-derived initiator gene.

However, this does not imply that replication initiation in the Fusobacteriaceae mirrors the process observed in species where DnaA drives chromosomal replication initiation. In support of this, the activation of iteron-plasmid Rep-type initiators is known to depend on the bacterial DnaK chaperone (the homolog of Hsp70 in eukaryotes) and its co-chaperones, DnaJ and GrpE. These initiators typically dimerize, yet they must bind to the replication origin as monomers; the chaperone system facilitates this monomerization and remodels the protein to promote binding (Wickner et al., 1991). Consequently, our work suggests that DnaK and its co-chaperones might be key players in Fusobacteriaceae replication initiation. To date, *Caulobacter crescentus* is the only major model organism for which DnaK is known to be directly involved in chromosomal replication initiation, with the notable exception of its role in initiating replication of the *Vibrio cholerae* chromosome 2, which is likely of iteron-plasmid origin (see Jha et al., 2017; Adam et al., 2022). Nevertheless, in *C. crescentus*, chaperones help accumulate sufficient levels of DnaA to allow replication initiation by restraining the synthesis of the protease Lon, which would otherwise degrade DnaA (Jonas *et al*., 2013). This involvement of DnaK in *C. crescentus* is thus fundamentally different from the mechanism hypothesized here for Fusobacteriaceae.

Identifying Replication initiator protein A (RPA) “partners” involved in Fusobacteriaceae replication initiation will require experimental validation. Nevertheless, from an evolutionary standpoint, our study provides the first documented case of a complete shift from a classical bacterial chromosomal replication system to one derived from a plasmid, underscoring the significant role of domesticated mobile elements in the evolution of the bacterial replication machinery (see (see Brézellec *et al*., 2006, 2016, 2017, 2026; Adam *et al*., 2023)).

While this study focused primarily on replication initiation, the contrast in oxygen tolerance between Leptotrichiaceae and Fusobacteriaceae (where the former are facultative or obligate anaerobes, whereas the latter range from microaerotolerant to obligately anaerobic (Gupta & Sethi, 2014)) suggests that the replication shift alone is insufficient to account for the full emergence of the Fusobacteriaceae family. A better understanding of this evolutionary event will thus require genome-wide comparisons between members of these two families. We speculate that our present findings, combined with future comparative genomics, may help shed light on *Fusobacterium nucleatum*, a member of the Fusobacteriaceae family known to be a driver of colorectal cancer progression (Udayasuryan *et al*., 2024). Specifically, the unconventional replication initiation mechanisms documented here, along with the possible involvement of the DnaK chaperone (typically active during exposure to environmental and physiological stress (Mayer, 2021)) might increase its fitness within the hypoxic tumor microenvironment, which represents a highly favorable niche for these bacteria.

## Supplementary information availability

Supplementary Tables S1 and S2 are available online as a gzip archive at https://doi.org/10.64898/2026.10.08.757545. Code and raw software outputs supporting this study are available on Zenodo at https://zenodo.org/records/23887865.

